# Improved analysis of CRISPR fitness screens and reduced off-target effects with the BAGEL2 gene essentiality classifier

**DOI:** 10.1101/2020.05.30.125526

**Authors:** Eiru Kim, Traver Hart

## Abstract

Identifying essential genes in genome-wide loss of function screens is a critical step in functional genomics and cancer target finding. We previously described the Bayesian Analysis of Gene Essentiality (BAGEL) algorithm for accurate classification of gene essentiality from short hairpin RNA and CRISPR/Cas9 genome wide genetic screens. Here, we introduce an updated version, BAGEL2, which employs an improved model that offers greater dynamic range of Bayes Factors, enabling detection of tumor suppressor genes, and a multi-target correction that reduces false positives from off-target CRISPR guide RNA. We also suggest a metric for screen quality at the replicate level and demonstrate how different algorithms handle lower-quality data in substantially different ways. BAGEL2 is written in Python 3 and source code, along with all supporting files, are available on github (https://github.com/hart-lab/bagel).

## Introduction

The landscape of preclinical studies to identify novel cancer targets has been fundamentally altered by the development of high throughput genome-wide CRISPR knock-out screens (1–3). The CRISPR-Cas9 system offers significant advantages in specificity and effectiveness of gene knock-out (3, 4) over the shRNA knock-down technology that preceded it. Genome-scale knockout screens enable the unbiased identification of genes that whose disruption impedes proliferation compared to wildtype cells (“essential genes”), and curation of pan- and context-dependent essential genes is being exploited to identify potential drug targets for specific tumor genotypes (3, 5–10). Precise analysis of genetic screen data is particularly important given recent evidence that off-target effects can mislead targeted drug development efforts (11).

Previously, we developed an effective algorithm, the Bayesian Analysis of Gene Essentiality (BAGEL), for classifying essential and nonessential genes in pooled library gene perturbation screens using either CRISPR or shRNA (12, 13). BAGEL calculates the log likelihood that a gene belongs to either the ‘essential’ or ‘nonessential’ class, and returns a log Bayes Factor (BF) that, in the context of a typical genome-scale knockout screen in a cell line, represents a blend of statistical confidence and biological effect size. The classifier is trained using gold-standard reference sets of likely core essential and nonessential genes, themselves derived from genetic screens and gene expression studies (12, 14). Provided appropriate care is taken to prevent circularity, these gold standards also offer an unbiased yardstick against which to compare the performance of other algorithms, screening technologies, and experimental designs.

Despite its utility, the previous version of BAGEL has some notable limitations. Firstly, the previous version used a truncated fold change model to calculate Bayes Factor, which capped the dynamic range of Bayes Factors. Secondly, though the bootstrapping approach it uses to train models is robust, it is computationally expensive, resulting in long run times under normal conditions. Lastly, there is no provision for correcting copy number amplification effects (5, 9, 15) or multi-targeting gRNA effects. To address these limitations, we have developed new version of the software, BAGEL2, which we present here. While the core algorithm remains intact, we present several changes that improve the runtime and accuracy of BAGEL, including a correction for gRNA off-target effects and an increased dynamic range of BFs that enables the detection of tumor suppressor genes whose knockout gives rise to increased cellular fitness. While BAGEL2 does not contain a method to address copy number artifacts, we describe a pipeline using CRISPRcleanR (16) for correcting these effects.

## Results

### An improved log likelihood/regression model

The analysis pipeline for a loss-of-function fitness screen consists of three steps: (1) mapping reads to the guide sequences in the CRISPR library and building a table of read counts, (2) normalizing counts across samples and calculating guide-level fold change, and (3) compiling guide-level information into gene-level fitness scores (Figure 1A). CRISPR screen analysis starts from the step of mapping raw sequencing read files to their corresponding CRISPR library. Mapping reads can be done with a variety of sequence analysis tools, including Bowtie (17), MAGeCK (18) and poolQ (https://portals.broadinstitute.org/gpp/public/software/poolq). Fold change is calculated by comparing endpoint to starting plasmid or T0 sample, a function now available in BAGEL2 using the *fc* option.

**Figure 1.**
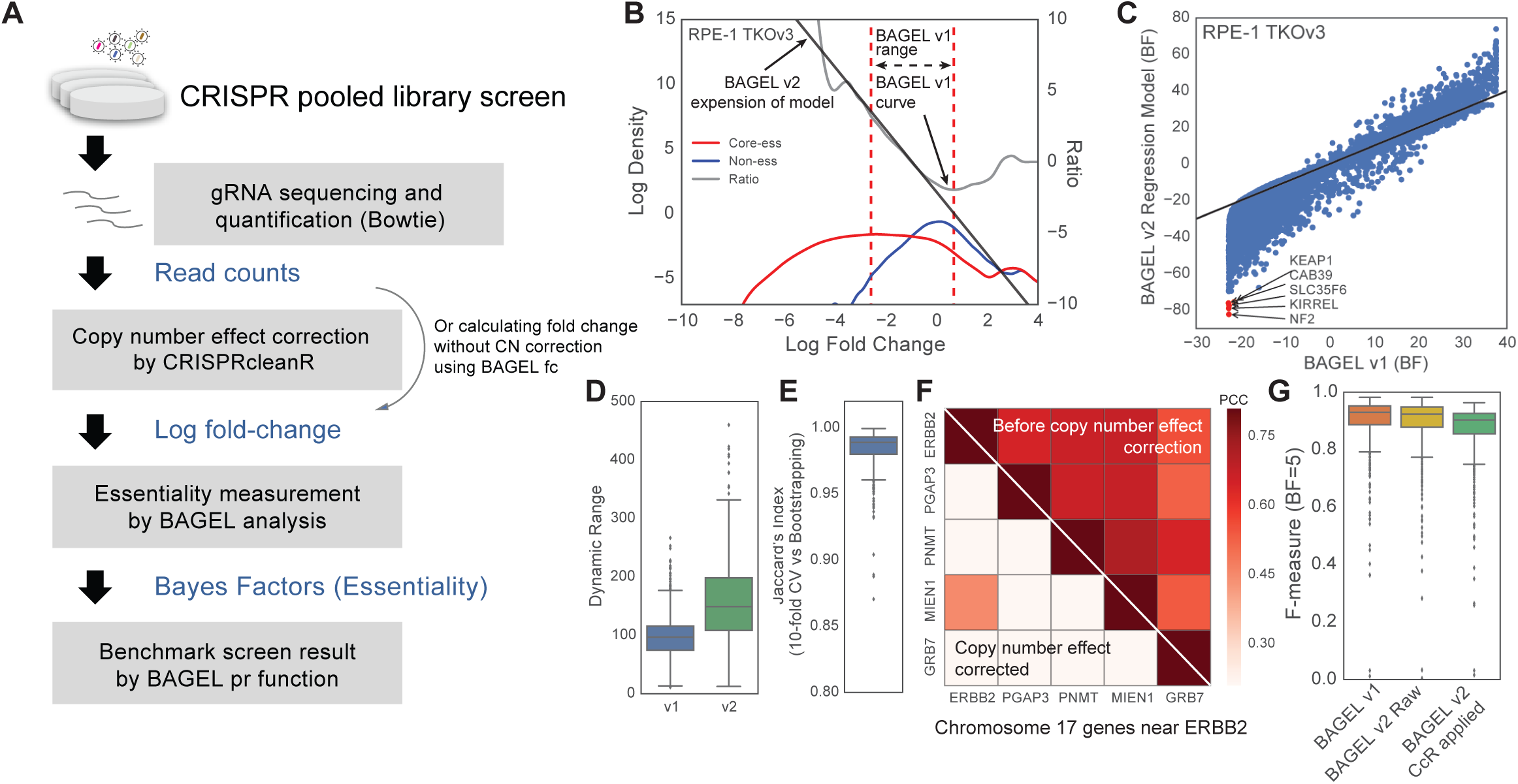
Improvement of BAGEL algorithm. **A)** A brief flow diagram of CRISPR pooled library screen analysis using BAGEL pipeline. **B)** Improved model of train model. The red and the blue curves indicate kernel density plots of fold changes of reference core-essential and non-essential genes, respectively. The grey curve indicates ratio of core-essential density to non-essential density at the point of fold change. Since there are a few data points in marginal area, BAGEL1 limited calculation area of fold change between the point that blue curve hits the density threshold (2-7 was used in BAGEL1) as a lower bound and the first local minimum ratio as an upper bound (red dashed lines). In BAGEL2, we employed linear regression to interpolate margin area in both ends (black line). **C)** Comparison of gene essentiality (Bayes Factor) between BAGEL1 and BAGEL2 using RPE-1 cell line screened by TKO v3. Known tumor suppressors (NF2, KIRREL, and KEAP1) that are scored BF ∼ −20 with hundreds of other genes in BAGEL1 were measured as much lower Bayes Factor and distinguished clearly with others in BAGEL2. **D)** Dynamic range of BAGEL2 results were increased from BAGEL1 across screens in Avana dataset. Jaccard index between predicted essential gene sets by 10-fold cross validation and bootstrapping. **F)** Pearson correlation coefficient of essentiality across 517 cell lines in Avana data between frequently amplified genes near ERBB2 on chromosome 17. After CRISPRcleanR applied, essentiality correlation due to copy number amplification effect was successfully corrected. **G)** Prediction performance benchmark between BAGEL1, BAGEL2 applied linear interpolation and 10-fold cross validation (BAGEL2 Raw), and BAGEL2 + CRISPRcleanR applied version (BAGEL2 CCR applied).

To calculate a gene essentiality score, BAGEL2 adopts the same Bayesian model selection approach as BAGEL. The “essential” model is represented by a kernel density estimate (KDE) of the distribution of guide-level fold changes of gRNA targeting a training set of essential genes (14), and the “nonessential” model is likewise trained on a set of nonessential genes (Figure 1B) (12, 14). Then, for each gRNA targeting each gene, a guide-level log Bayes Factor (BF) is calculated as the log ratio of these two kernel density estimates, evaluated at the observed log fold change of the guide.

The stability of this calculation depends heavily on the local density of data points used to calculate the training set KDEs. For example, at extreme fold changes, sparsity of training data from the nonessential set results in extreme ratios. For this reason, in the previous version of BAGEL, we defined the boundaries of the near-linear rage of this ratio and truncated all data outside these boundaries (Figure 1B). Guide-level log BFs are then summed to gene-level log BFs (hereafter all Bayes Factors are in log2 space). BAGEL2 relaxes this limitation by calculating a linear best fit to the log ratio in this space and using this fit to extrapolate the BF calculation to all observed fold changes (Figure 1B, gray line). The net result is a better usage of the total fold change data, a correction for log-ratio instability at positive or extreme negative guide-level fold changes, and a broader dynamic range of gene-level BFs reported by the algorithm.

An unanticipated result of this broader dynamic range is that BAGEL2 now detects candidate tumor suppressor genes. We re-analyzed a previously reported genome-scale screen of RPE1 retinal pigmented epithelium cells performed with the TKOv3 library (14), and comparing BAGEL to BAGEL2 results shows the truncation of gene-level BFs in BAGEL (Figure 1C). Notably, outliers with extreme negative BFs (Figure 1C, red) include genes with known tumor suppressor activity, including *KEAP1* (19, 20) and Hippo pathway genes *NF2* and *KIRREL* (21). We confirmed the regression scheme increases dynamic range through hundreds of cell lines in Avana dataset downloaded from DepMap (5) **(Figure 1D)**.

Another improvement in BAGEL2 involves replacing bootstrapping with 10-fold cross-validation. Bootstrap resampling of the training sets, used in BAGEL, provides a robust method to evaluate the effect of training data variance on gene-level BF calculations, but is computationally expensive. However, given the large size of training sets used for genome-scale fitness screens, sampling introduces relatively little variance. Ten-fold cross validation yields nearly identical Bayes Factor distributions as bootstrapping in most cases, and comparing BAGEL and BAGEL2 hits (BF>=5) in the DepMap data yields Jaccard coefficients ∼0.99 (**Figure 1E**). Cross-validation is the default setting in BAGEL2, and speeds up running time on a single processor by ∼50-fold.

Copy number amplifications are a known source of potential artifacts in CRISPR knockout fitness screens (9, 15, 22), and BAGEL2 does not correct for this source of error. Instead, we employed an unsupervised copy number correction algorithm, CRISPRcleanR (16), as a preprocessing step. CRISPRcleanR corrects amplicon-induced artifacts based on guide position and fold change, without copy number information. We find that BAGEL2 with copy number correction preprocessing successfully reduces amplicon-induced artifacts (Figure 1F) while maintaining high sensitivity and specificity (Figure 1G). Overall, BAGEL2 improves performance and sensitivity over BAGEL at no additional cost.

### Correcting Multi-targeting effects and false positive analysis

It is widely accepted that the specificity and sensitivity of CRISPR reagents far exceeds that of prior-generation shRNA reagents (4). However, off-target effects of CRISPR reagents can still confound loss-of-function screens. Recently, several studies reported that CRISPR/Cas9 reagents have a non-negligible effect on off-target cut sites with mismatches of 1-2bp from the intended target site (23, 24). These off-targeting effects by mismatched targets can cause additional ad hoc DNA cutting or, depending on the locus, knockout of genes. We found that many of guide RNAs in the Avana and KY libraries that target several sites with perfect matches, and our TKOv3 library was specifically designed to allow up to one perfect-match, off-target cut site in an intragenic region (14) (Supplementary Figure 1). These multi-targeting gRNAs can result in unexpected fitness defect, the effect of which can be decomposed into target-specific and off-target/multiple-targeting effects (See Methods). To implement multi-targeting effect correction in a single cell line screen, we re-aligned gRNA sequences of CRISPR/Cas9 libraries to the human genome with mismatches allowed. In this study, we only consider perfect matched targets and 1-bp mismatched targets. Using alignment information, BAGEL discards promiscuous gRNAs (perfect match > 10 loci or 1-bp mismatch > 10 loci) from libraries. Then, to measure the component of fitness defect specific to multi-targeting effects, we took sgRNAs targeting multiple loci with 0-1 mismatches to non-protein-coding regions (excluding protein-coding off-target sites to minimize the contribution from genetic interactions; Supplementary Figure 1). The multiple-targeting effects of gRNAs can be estimated by the incremental BF in comparison with gRNAs targeting the same gene but with no off-target cut sites. For example, in ovarian endometrioid cancer cell line OVK-18, the multiple-targeting effects of the Avana library showed an incremental BF due to off-targets that increased roughly linearly with the number of perfect-match, off-target cut sites in the genome and a smaller incremental guide-level BF with the frequency of mismatched off-target sites (Figure 3A; Supplementary Figure 1B). Since we only addressed sgRNAs targeting multiple loci but targeting only one protein coding locus, these effects were exclusively from multi-targeting effects, not the effect of genetic interaction. In our example case, each additional perfect-match target boosted the BF of a single gRNA by 3.5 and each additional 1bp-mismatched target increased the BF by 1.4 (∼40% of the boost attributable to perfect matches). We removed these off-target effects by guide-level regression of incremental BF vs. off-target effects (i.e. applied a BF penalty based on the number of predicted off-target cut sites) and confirmed that the bias was no longer present after the effect was removed (Figure 2B).

**Figure 2.**
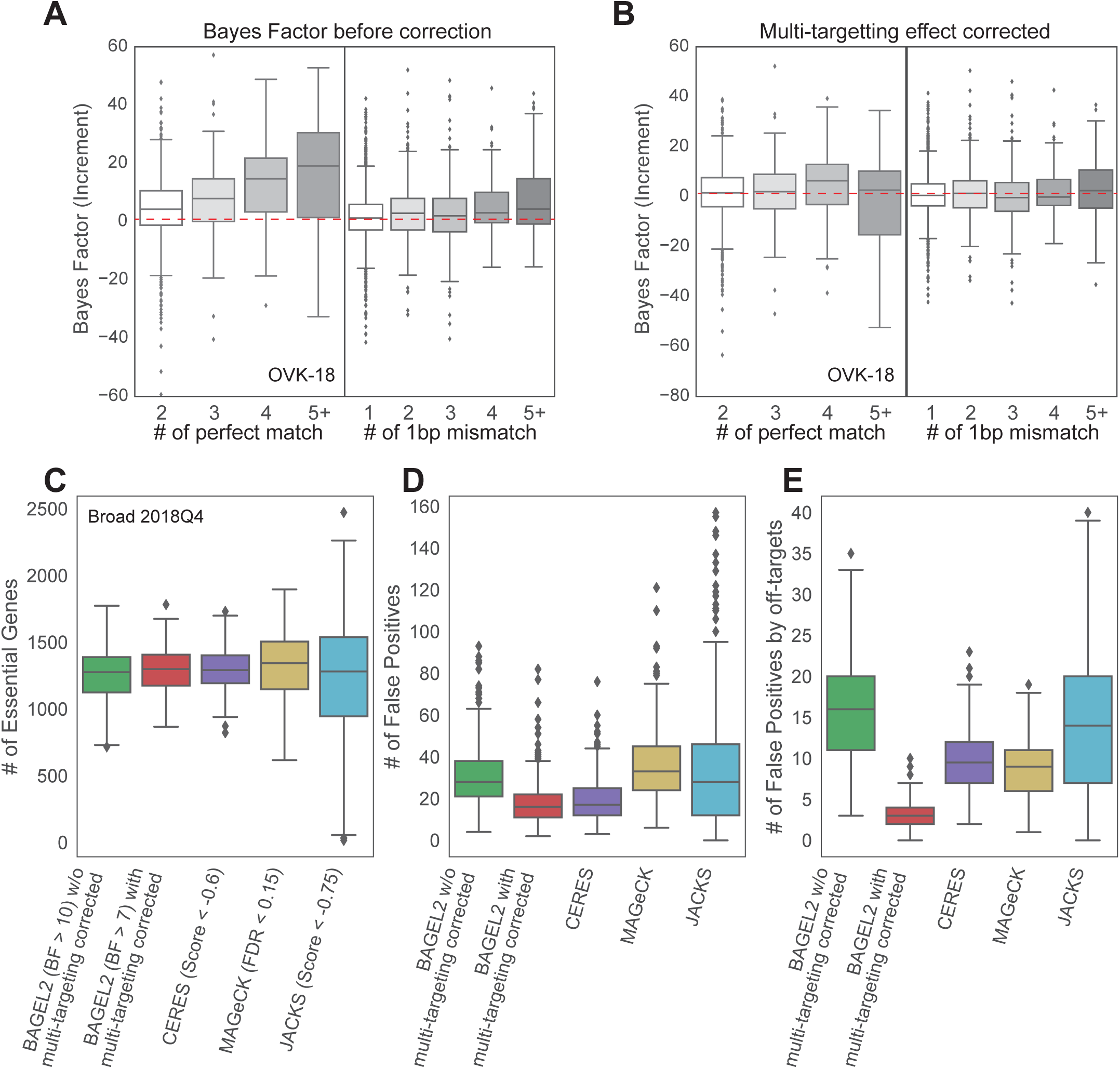
The multi-targeting effect correction reduces false positives from off-targets with 1-bp mismatch. **A,B)** Increment of Bayes Factors of multi-targeting gRNAs but targeting only single protein coding gene in comparison with Bayes Factor of gRNAs targeting the protein coding gene without any other targets **A)** before the multi-targeting effect correction and **B)** after the multi-targeting effect correction. **C)** The targeting effects from perfect-matched targets and 1bp-mismatched targets was well correlated across 517 cell lines in Avana dataset. **D)** The number of essential genes across good quality cell lines (F-measure > 0.85) in Avana dataset predicted by BAGEL2 and other algorithms, CERES, MAGeCK, and JACKS with cut-off threshold BF 10, score –0.6, FDR 0.15, and score –0.6, respectively. The cut-off threshold was aimed for obtaining similar numbers of essential genes. **E)** The number false positives predicted by each algorithm. False positives were defined by non-expressed genes in RNA-seq data of corresponding cell lines. BAGEL2 after multi-targeting effect correction shows comparable results with CERES and much lower numbers than results of MAGeCK and JACKS. **F)** The number of false positives in predicted essential genesets when the scope is limited to genes having gRNAs mapped over than five 1-bp mismatched targets that are likely from multi-targeting effects of 1-bp mismatched targets. The result of BAGEL2 after correction shows the best performance among algorithms.

**Figure 3.**
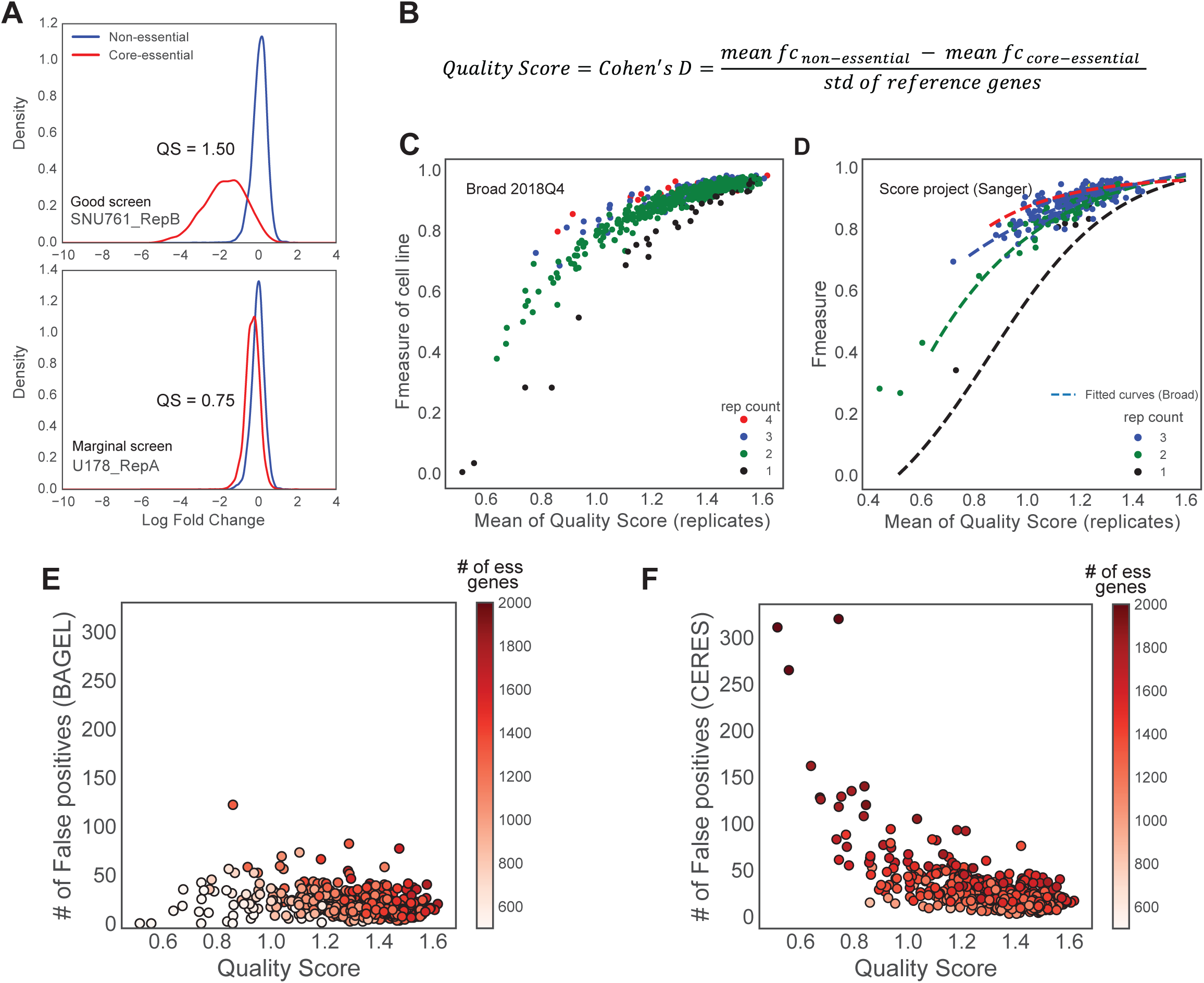
the good screen and the bad screen. **A)** The kernel density curves of reference core-essential genes (red) and non-essential genes (blue) with SNU761 replicate B as an example of good screen (upper panel) and U178 replicate A as an example of marginal screen (lower panel). The good screen shows clear separation between core-essential and non-essential curves whereas the marginal screen shows less separation. **B)** The equation of quality score. **C)** Mean quality score of replicates and f-measure in cell line level shows clear correlation trends and differentiated by replicate screen counts per cell line. **D)** Down-sampling analysis of replicate counts using Achilles CRISPR screen data recapitulated and followed same trends of Avana set. **E**, A plot of relationship between the number of false positives in **E)** BAGEL results and **F)** CERES results across 517 cell lines in Avana data. Each dot colored by the number of essential genes.

We compared BAGEL2 with multi-targeting correction to BAGEL2 without correction, as well as to other contemporary screen analysis algorithms, including CERES, MAGeCK, and JACKS, run against the DepMap 2018Q4 data release, using raw read counts per guide as a starting point. Since the number of false positives is sensitive to the number of essential genes, we identified thresholds for each algorithm that returned roughly the same median number of essential genes across the 518 cell line screens analyzed. We used cut-off values of BF 10, BF 7, score −0.6, FDR 0.15, and score −0.75 for BAGEL2 uncorrected, BAGEL2 with multitargeting effect correction, CERES (5), MAGeCK (18), and JACKS (25), respectively (Figure 2C). Then, for each cell line, we identified a set of genes with no to low expression (logTPM < 1), judging that genes with trace mRNA expression levels cannot be essential and following the concept used to define the non-essential reference gene set (12). For each algorithm, we identified the total number of expression-defined false positives (Figure 2D). BAGEL2 results after multi-targeting effect correction showed the lowest number of false positives and resulted significantly lower false positives than MAGeCK and JACKS, while CERES showed a similar number of false positives with BAGEL2. To further investigate whether the correction can minimize false positives from multi-targeting effects, we limited the scope to non-expressed genes targeted by gRNA with 5 or more 1bp-mismatched off-target cut sites in the genome. BAGEL2 multitarget correction effectively filters these genes. (**Figure 2E**). We also demonstrate that agreement of gene essentiality across cell lines screened using both the Avana and KY libraries can be improved by multi-targeting effect correction (**Supplementary Figure 2**). Overall, we demonstrated new version of BAGEL can correct the multi-targeting effects from perfect-matched and 1bp-mismatched targets, reducing the number of false positives arising from multi-targeting effects. Also, we showed that BAGEL algorithm identified essential genes accurately in comparison with other algorithms through false positive analysis.

### Replicate quality score can predict performance of cell-line screens

CRISPR screens require significant technical expertise, but even in the best hands results can vary for numerous reasons, including environmental, experimental, and intrinsic factors such as batch effect, PCR noise, stochastic off-target events of guide RNAs, and characteristics of individual cell lines (24, 26–28). Understanding and identifying effective and ineffective screens is necessary to understanding gene essentiality and differential essentiality. Previously, we defined lists of core-essential and non-essential genes (12, 14). These reference gene sets are not only used as training sets in BAGEL, but also can be used to evaluate the quality of a screen (**Figure 3A**). We compared a good screen (SNU-761, replicate B) and a marginal screen (U-178, replicate A) from the same batch in the Avana dataset. While the good screen shows clear separation of core-essential (red) and non-essential guides (blue), the marginal screen shows much greater overlap between the two distributions. To distinguish good from marginal screens, we employed as a quality score the Cohen’s D statistic, which is the difference in mean log fold change between core-essential and non-essential genes divided by the standard deviation of log fold change of all reference genes (**Figure 3B**). The maximum possible quality score is 2.0. Using this scheme, we calculated the quality scores of SNU-761 (rep B) and U-178 (rep A) as 1.50 and 0.75, respectively (Figure 3A).

We further applied this quality score measurement to all individual replicates in Avana dataset, and we compared with the F-measure of cell lines that is derived by BAGEL2 aggregation of all replicates (**Figure 3C**). Many low-performing cell lines (F-measure < 0.7) were from replicates having low mean quality score (below 1.0). Notably, even when average replicate quality is less than ideal, multiple replicates can boost overall screen quality; likewise, even high-quality, single-replicate screens show lower overall F-measure than equivalent quality screens with two or more replicates. We confirmed the generality of these trends by conducting the same analysis with data from Project Score (6) (**Figure 3D**). The relationship between replicate quality score, number of replicates per screen, and overall screen F-measure was highly consistent with the Broad data, and supports the applicability of the Cohen’s D statistic as a replicate-level quality score.

Although quality scores of replicates were directly related to the overall reliability of an experiment, we evaluated the effect of including one low-quality replicate in an otherwise high-quality screen. Although the overall trend supports the general notion that additional replicates can smooth out random noise and increase overall screen performance (**Figure 3C,D**), we find that a single low-quality replicate (Quality score < 1.0) among one or more high confident replicates (Quality score > 1.4) can reduce overall performance (**Supplementary Figure 4**).

### Data quality has different effects on different algorithms

The quality of the underlying data can have profound effects on the results of an analytical pipeline. We compared how quality score affect the results of BAGEL2 and CERES (**Figure 3E, 3F, Supplementary Table 1**). Interestingly, the two algorithms show opposite behavior as the quality of the underlying data degrades. In BAGEL2, the number of false positives remained a similar across all quality levels, but BAGEL2 calls very few essential genes for low-quality data (**Figure 3E**). In contrast, CERES amplifies the number of hits and the corresponding number of false positives as quality degrades (**Figure 3F**). These results reflect the approaches adopted by the two algorithms. The Bayes Factor approach provides a summary statistic that essentially combines effect size and statistical significance. Since lower quality screens offer both lower effect size (fold change) and corresponding statistical power, the number of essential genes in a lower quality screen will be fewer than in a high-quality screen. On the other hand, CERES rescales results by setting the score of core essential genes to −1.0 and non-essential genes to zero. Since low quality screens poorly distinguish between essential genes and non-essential genes (**Figure 3A**), significant error can be introduced by this rescaling. It should be noted that most CRISPR data in the DepMap and Project Score are of sufficiently high quality that this is not an important factor (92% of screens have quality scores > 1.0); nevertheless, researchers should be wary when including marginal quality screens in their analyses.

## Discussion

In this study, we introduced an improved version of BAGEL algorithm, BAGEL2, for genome-wide pooled-library loss-of-function fitness screens. We showed the linear interpolation of score expands the dynamic range of Bayes Factor in comparison of previous version of BAGEL, enabling more accurate quantitation of fitness defects as well as discovery of putative tumor suppressor genes whose knockout results in faster proliferation. Also, we demonstrated that correction of multiple target effect can successfully remove false positives.

We show that BAGEL2 can remedy false positives caused by CRISPR multi-targeting guides. That these effects can be mitigated algorithmically is important and useful. However, in the future this effect should be addressed at the library design level. In particular the Avana library contains many multi-targeting sgRNAs, compared to the Brunello (29), TKOv3 (14), and KY libraries (6). However, there is clearly an advantage to screening with the Avana library and comparing results with the large and growing corpus of cell line characterization data available. Researchers will have to make their own informed decisions weighing these advantages and disadvantages.

We suggest Cohen’s D statistic, evaluated against core-essential genes and non-essential genes, to provide a quantitative measure of the quality of single screen replicates. We show that, as expected, the number and quality of replicates is directly related to the overall screen performance (F-measure). Interestingly, however, we also show that individual “bad” replicates can degrade the overall performance of an otherwise “good” screen. Therefore, we recommend evaluating quality at the replicate level and discarding low-quality outliers from groups of otherwise high-quality replicates.

The software can be downloaded through github (https://github.com/hart-lab/bagel).

## Materials and Methods

### BAGEL pipeline summary

As an input of BAGEL pipeline, a tab separated plain text of read count file can be used. We employed external application, CRISPRcleanR (16), to calculate fold change with copy number effect correction. In addition, there is a built-in ‘fc’ function in BAGEL application to calculate fold change who don’t need to correct copy number effect correction. After that, essentiality was calculated using BAGEL ‘bf’ function from the fold change file. Finally, benchmarking by precision and recall of reference genes were conducted using BAGEL ‘pr’ function.

### Preparing a read count file

If it starts from a fastq file of reads, alignment into reference gRNA library can be conducted by Bowtie version 1.1.2. For best accuracy, parameters -v 0 -m 1 to search perfect matched read only (-v 0), limit results maximum 1 (-m 1) was used. Then, read counts can be generated by parsing the result sam file.

### Calculate fold change from read count file and correct copy number effect using CRISPRcleanR

CRISPRcleanR was downloaded from github (https://github.com/francescojm/CRISPRcleanR). To run CRISPRcleanR, we built alignment information of a CRISPR library. We mapped position and exon of gRNAs using gencode annotation v28 for GRCh37. Since CRISPRcleanR generated one summarized fold changes for all input replicates, we ran CRISPRcleanR for each replicate separately. Then, pasted them into one file as separate columns. Otherwise this, we ran CRISPRcleanR by the default practice provided by author.

### Bayes Factor (BF) calculation – BAGEL ‘bf’ function

BAGEL ‘bf’ function is a tool for calculating Bayes Factor, which is a degree of essentiality (12). The formula of Bayes Factor defined previously is as below.

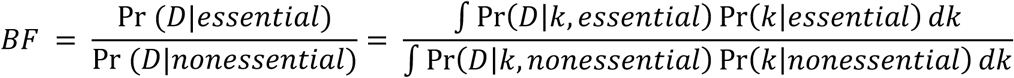

To implement this, BAGEL samples all genes into training set and test set by user defined methods between 10-fold cross validation or bootstrapping. In each iteration, BAGEL build linear regression model using log density ratio values of core-essential to non-essential for every fold change value between the fold change where the density of non-essential curves meets predefined threshold and the fold change where the log density ratio is the lowest (**Figure 1B**). Core-essential and non-essential gene set are reference gene lists defined in previous study (12, 13). Guide RNA level Bayes Factor is derived by the sum of the linear regression model results by fold changes of each replicate. And, gene level Bayes Factor is calculated by sum of all gRNA level BFs mapped to the gene.

### Correcting multi-targeting effects

Essentiality of guide RNA is the sum of the effects from target gene, DNA and genetic interaction between target genes.

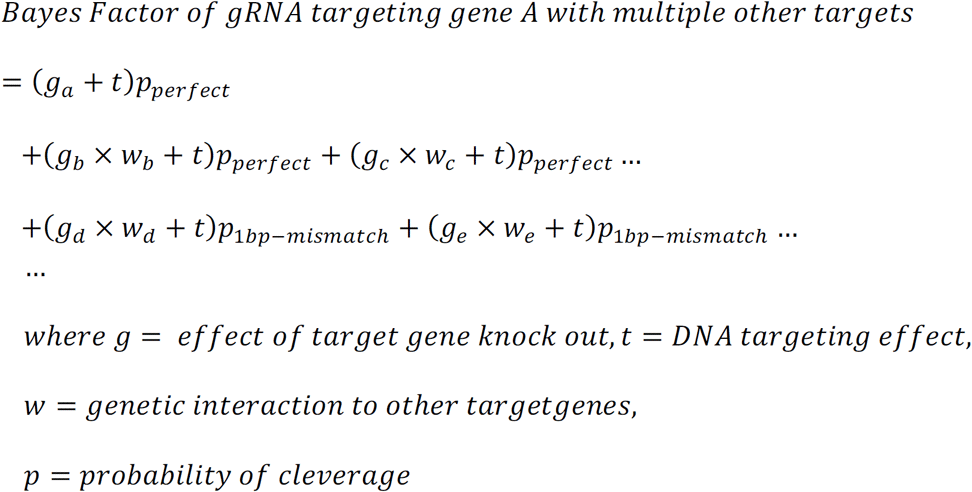

Since genetic interaction effect is difficult to deconvoluted from gene effects, we only correct DNA targeting effect from multiple DNA regions. In this formula, we supposed to measure DNA targeting effect of perfect matched targets and 1bp-mismatched targets. So, we collected data points from gRNAs targeting multiple non-coding regions with perfect match or 1bp mismatch except one desired protein-coding gene. Since the regions except one desired protein coding locus don’t be affected by gene knock out effect, the formula is like below.

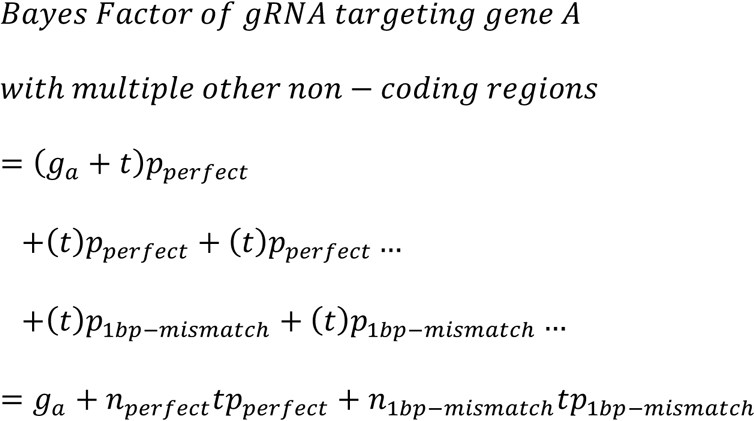

Then, we inferred *tp*_*perfect*_ and *tp*_1*bp* − *mismatch*_ by multiple linear regression (**Figure 2A, 2B**, and **Supplementary Figure 1**). Finally, we gave penalties Bayes Factor of gRNAs based on the number of perfect targets and 1bp-mismatched targets.

### F-measure and false discovery rate (FDR) calculation

F-measure (BF=5) is a harmonic mean of precision and recall at the threshold of Bayes Factor 5 and it can represent performance of essentiality prediction. We calculate f-measure using a pr table generated by BAGEL pr function.

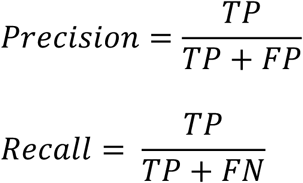

TP = positives in reference core-essential set

FP = positives in reference non-essential set

FN = negatives in reference core-essential set

Since it’s rare that a gene is exactly BF 5, we used the precision and the recall that is the nearest but above than BF 5. False discovery rate used in this study was calculated by 1.0 – precision.

### Acquirement of publicly available screen

There are large screen dataset providing CRISPR screens for cancer cell lines publicly such as Depmap project (Avana dataset) by Broad institute (5) and Score project by Sanger institute (Score dataset) (6). We downloaded read count data of Avana 2018Q4 dataset, which contains screens of 517 cancer cell lines, from the Depmap official website (http://www.depmap.org). Since Avana library allows targeting multiple protein coding genes, we discarded gRNAs targeting multiple protein coding genes without mismatch at the read count level data. We also downloaded read count data of Score project from the official website (https://depmap.sanger.ac.uk/). Then, we applied standard BAGEL2 pipeline with CRISPRcleanR copy number effect correction.

*Essentiality calculation using other dependency identifiers, MAGeCK, JACKS, and CERES* We downloaded MAGeCK (18) version 0.5.5 from the MAGeCK distribution website (https://sourceforge.net/p/mageck/wiki/Home/) and ran through Avana read count data with default parameters. For CERES (5), we used pre-calculated 2018Q4 dependency data downloaded from Depmap official website. We also downloaded JACKS (25) from the official github page (https://github.com/felicityallen/JACKS) and ran for Avana read count data with gene guide map and replicate information. To decide whether a gene is essential or not, we used ‘neg|fdr’ for MAGeCK and dependency score for CERES and JACKS.

### False positive analysis

To measure false positives of screen, we downloaded CCLE RNA-seq log TPM data for ∼1000 cell lines from DepMap (30). False positives of each cell lines were defined log-expression below than 1.0. Since the number of false positives were sensitive to the number of essential genes, we used thresholds of essential genes to keep similar level of number (**Figure 2C**). The thresholds were BF 10 for BAGEL2 data without correction and BF 7 for BAGEL2 data with multi-targeting correction. And, we used score −0.6, FDR 0.15, and score −0.75 for CERES, and MAGeCK, and JACKS, respectively. Then, we counted how many false positives are existed in essential gene sets defined by each algorithm across good quality cell lines (F-measure > 0.85) in Avana dataset.

### Quality score analysis

Measuring quality score of single replicate is important to decide experiment strategy. We measured Cohen’s D of reference core-essential genes and non-essential genes as a quality score of single replicates.

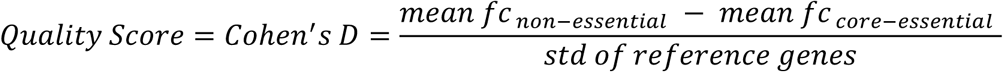

For each single replicate in Avana dataset, we collected all log fold changes values of gRNAs targeting either reference core-essential genes or non-essential genes. To demonstrate relationship between prediction performance and quality of single replicate screen, we compared F-measure (BF=5) from Bayes Factor using all replicates and mean quality score of each replicates calculated from fold change level. Down-sampling simulation using Achilles CRISPR data were conducted by picking replicates randomly in different numbers (single replicates ∼ 4 replicates) and calculated F-measure of them.

## Supplementary Table

Supplementary Table 1. The number of dependency calls in BAGEL2 (BF>7) and CERES (score < −0.6) and false positives in the calls across DepMap 2018Q4 screens with expression data (515 cells).

## Supplementary Figures

**Supplementary Figure 1.**
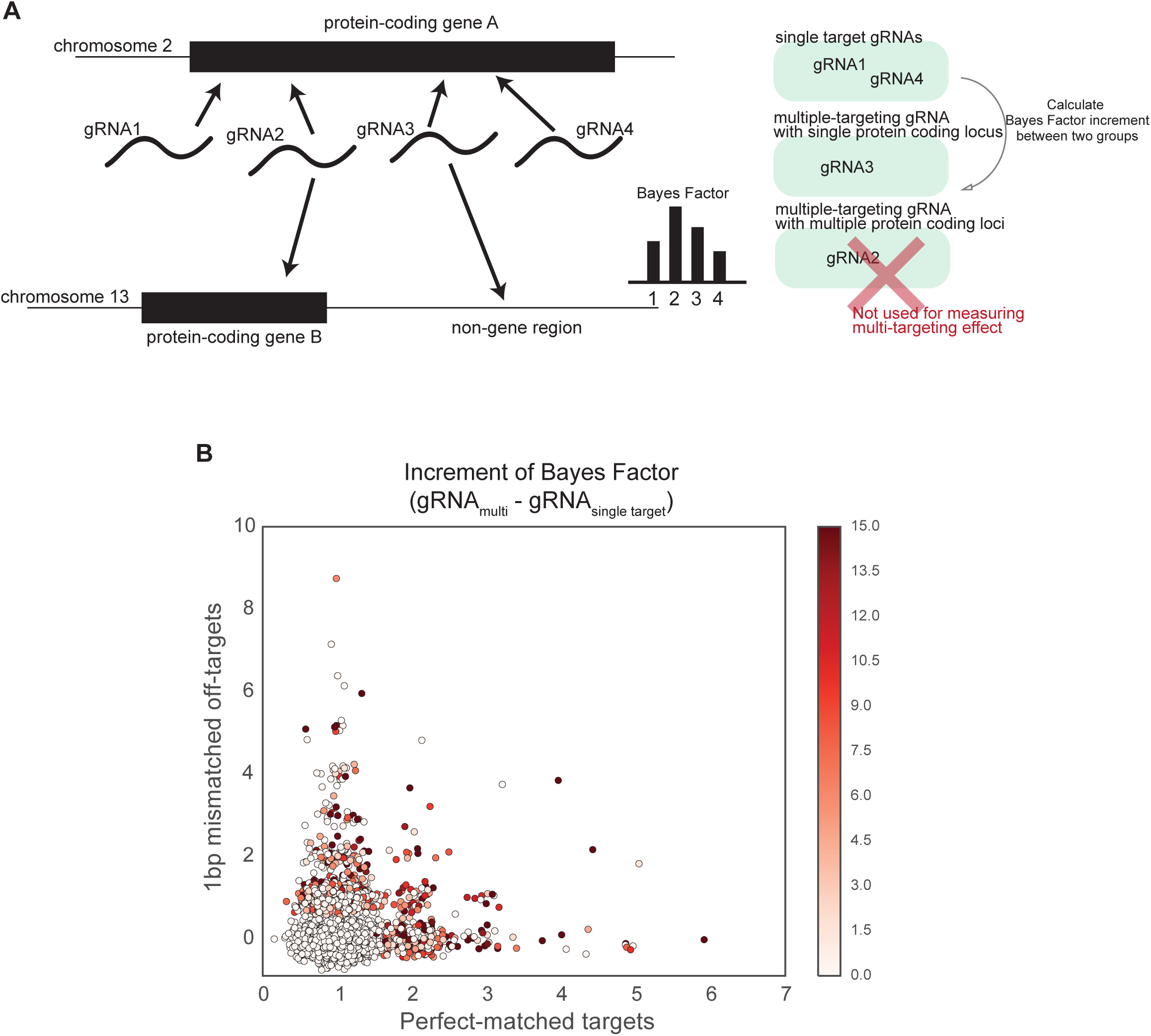
**A)** A plot explains how to calculate increment of Bayes Factor for measuring multi-targeting effect. **B)** A two-dimensional dot plot gRNAs targeting multiple regions but only targeting one protein-coding gene. Each dot is located at the number of perfect-matched targets and 1-bp mismatched targets with random jitter and colored by increment of Bayes Factor.

**Supplementary Figure 2.**
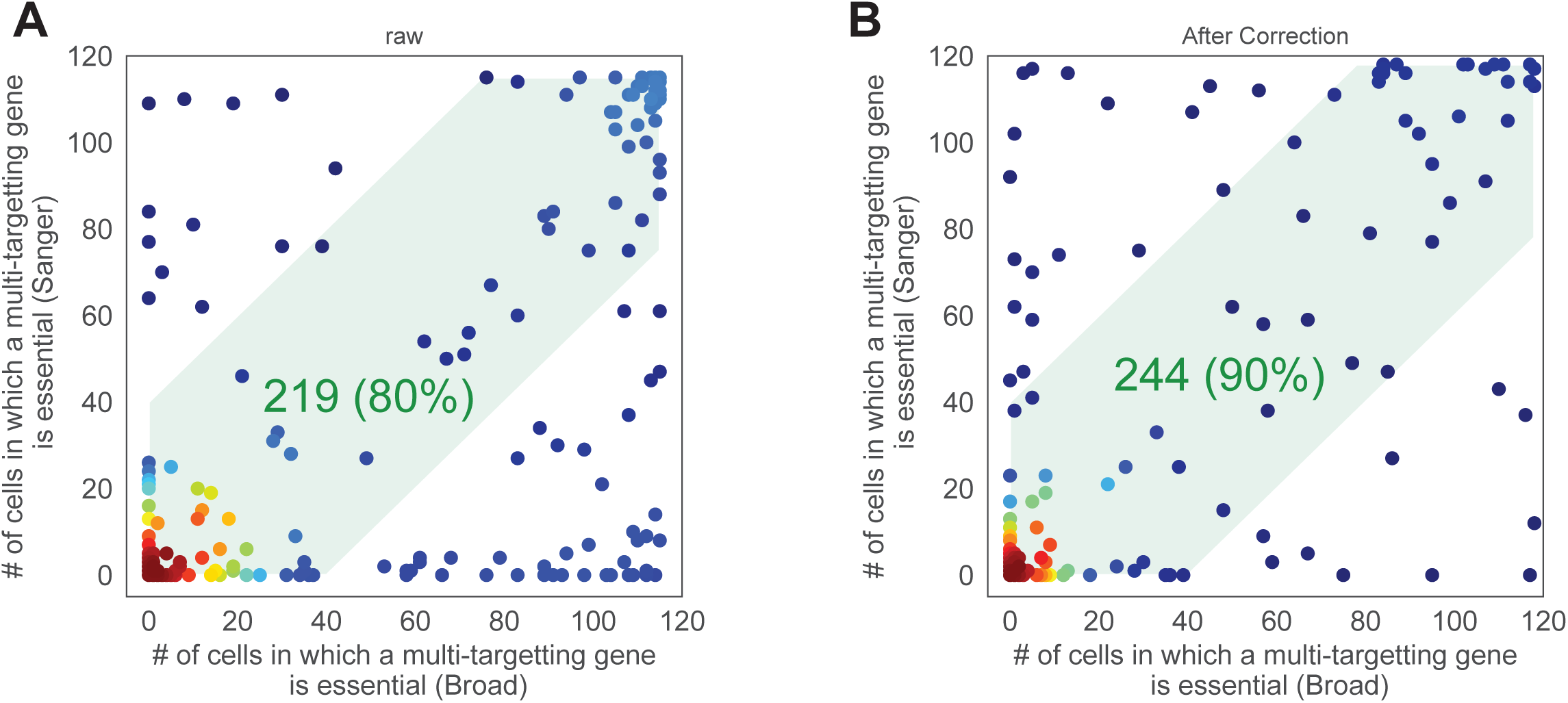
**A,B)** Agreement of genes that have gRNAs targeting 5 of more regions with 1-bp mismatch between Sanger data (Score project data) and Broad data (Avana dataset) **A)** before multi-targeting effect correction and **B)** after multi-targeting effect correction.

**Supplementary Figure 3.**
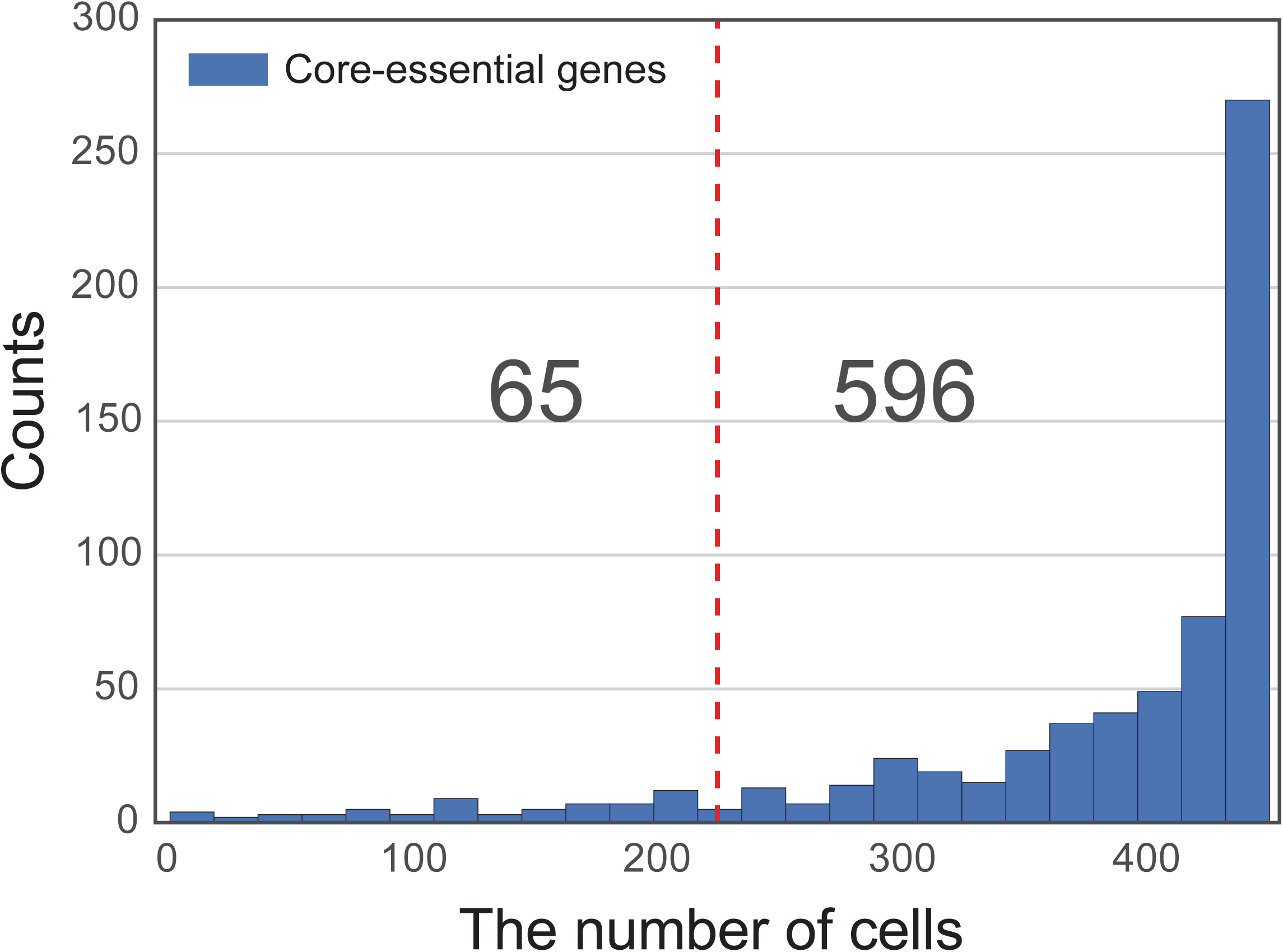
A histogram of the number of cells where genes are essential (BF > 5). The vertical dashed line indicates 50% of total cells.

**Supplementary Figure 4.**
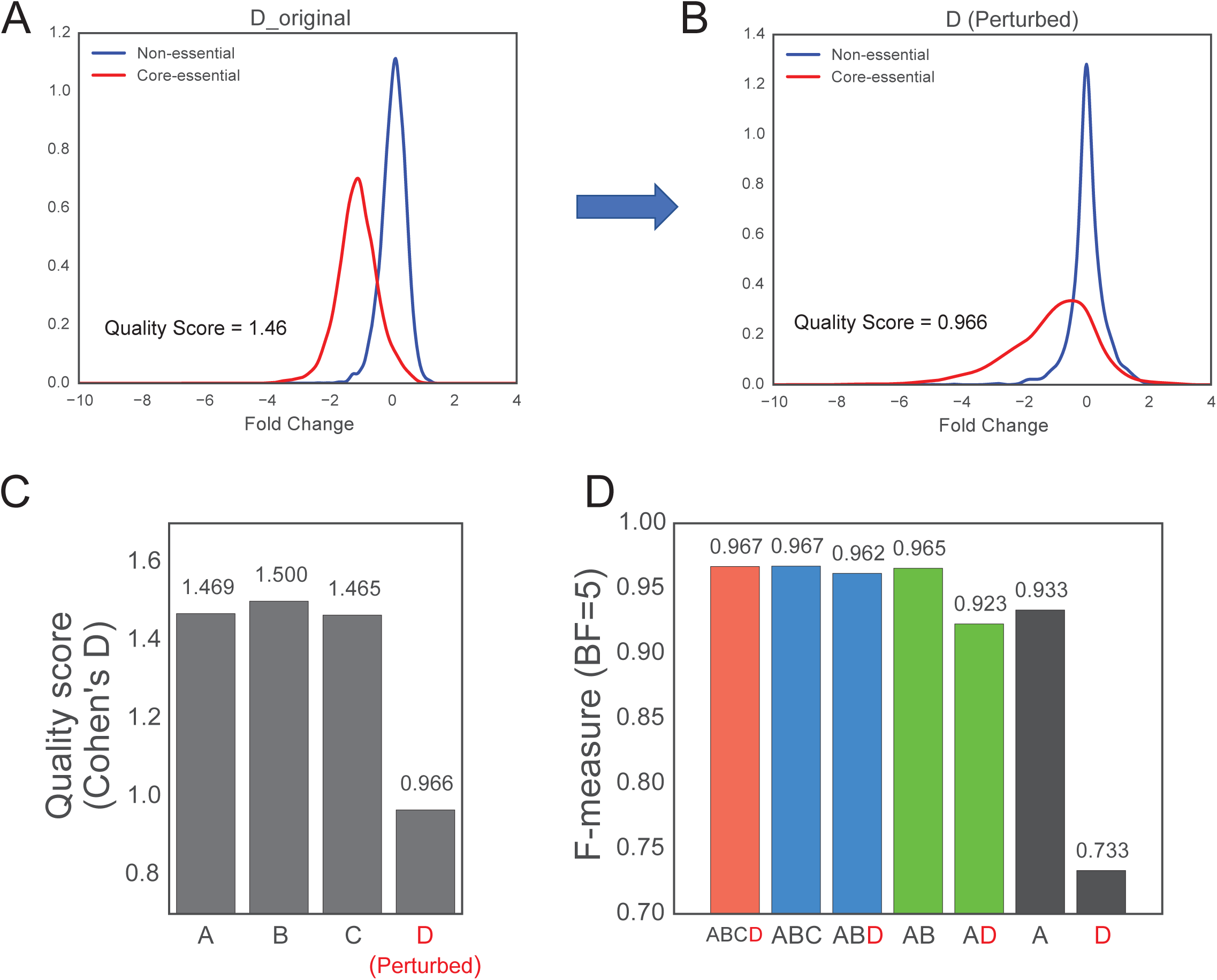
**A,B)** Fold change distribution plots for a replicate of HUP-T3 cell (A) before and (B) after fold change perturbation. To generate an low performance outlier sample, we added random noise to foldchange value. **C)** Quality scores of each replicate. **D)** F-measures (BF = 5) of combination of replicates. Adding the outlier (replicate D) to other high confident replicates reduce overall performance in condition of a few replicates (A vs AD and AB vs ABD).

## References

1. Zhou, Y., Zhu, S., Cai, C., Yuan, P., Li, C., Huang, Y. and Wei, W. (2014) High-throughput screening of a CRISPR/Cas9 library for functional genomics in human cells. Nature, 509, 487–491.

2. Shalem, O., Sanjana, N.E., Hartenian, E., Shi, X., Scott, D.A., Mikkelsen, T.S., Heckl, D., Ebert, B.L., Root, D.E., Doench, J.G., et al. (2014) Genome-Scale CRISPR-Cas9 Knockout Screening in Human Cells. Science, 343, 84–87.

3. Hart, T., Chandrashekhar, M., Aregger, M., Steinhart, Z., Brown, K.R., MacLeod, G., Mis, M., Zimmermann, M., Fradet-Turcotte, A., Sun, S., et al. (2015) High-Resolution CRISPR Screens Reveal Fitness Genes and Genotype-Specific Cancer Liabilities. Cell, 163, 1515–1526.

4. Evers, B., Jastrzebski, K., Heijmans, J.P.M., Grernrum, W., Beijersbergen, R.L. and Bernards, R. (2016) CRISPR knockout screening outperforms shRNA and CRISPRi in identifying essential genes. Nat. Biotechnol., 34, 631–633.

5. Meyers, R.M., Bryan, J.G., McFarland, J.M., Weir, B.A., Sizemore, A.E., Xu, H., Dharia, N.V., Montgomery, P.G., Cowley, G.S., Pantel, S., et al. (2017) Computational correction of copy number effect improves specificity of CRISPR-Cas9 essentiality screens in cancer cells. Nat. Genet., 49, 1779–1784.

6. Behan, F.M., Iorio, F., Picco, G., Gonçalves, E., Beaver, C.M., Migliardi, G., Santos, R., Rao, Y., Sassi, F., Pinnelli, M., et al. (2019) Prioritization of cancer therapeutic targets using CRISPR-Cas9 screens. Nature, 568, 511–516.

7. Wang, T., Birsoy, K., Hughes, N.W., Krupczak, K.M., Post, Y., Wei, J.J., Lander, E.S. and Sabatini, D.M. (2015) Identification and characterization of essential genes in the human genome. Science, 350, 1096–1101.

8. Wang, T., Yu, H., Hughes, N.W., Liu, B., Kendirli, A., Klein, K., Chen, W.W., Lander, E.S. and Sabatini, D.M. (2017) Gene Essentiality Profiling Reveals Gene Networks and Synthetic Lethal Interactions with Oncogenic Ras. Cell, 168, 890-903.e15.

9. Aguirre, A.J., Meyers, R.M., Weir, B.A., Vazquez, F., Zhang, C.-Z., Ben-David, U., Cook, A., Ha, G., Harrington, W.F., Doshi, M.B., et al. (2016) Genomic Copy Number Dictates a Gene-Independent Cell Response to CRISPR/Cas9 Targeting. Cancer Discov, 6, 914–929.

10. Steinhart, Z., Pavlovic, Z., Chandrashekhar, M., Hart, T., Wang, X., Zhang, X., Robitaille, M., Brown, K.R., Jaksani, S., Overmeer, R., et al. (2017) Genome-wide CRISPR screens reveal a Wnt–FZD5 signaling circuit as a druggable vulnerability of RNF43 -mutant pancreatic tumors. Nat Med, 23, 60–68.

11. Lin, A., Giuliano, C.J., Palladino, A., John, K.M., Abramowicz, C., Yuan, M.L., Sausville, E.L., Lukow, D.A., Liu, L., Chait, A.R., et al. (2019) Off-target toxicity is a common mechanism of action of cancer drugs undergoing clinical trials. Science Translational Medicine, 11.

12. Hart, T., Brown, K.R., Sircoulomb, F., Rottapel, R. and Moffat, J. (2014) Measuring error rates in genomic perturbation screens: gold standards for human functional genomics. Molecular Systems Biology, 10, 733.

13. Hart, T. and Moffat, J. (2016) BAGEL: a computational framework for identifying essential genes from pooled library screens. BMC Bioinformatics, 17, 164.

14. Hart, T., Tong, A.H.Y., Chan, K., Van Leeuwen, J., Seetharaman, A., Aregger, M., Chandrashekhar, M., Hustedt, N., Seth, S., Noonan, A., et al. (2017) Evaluation and Design of Genome-Wide CRISPR/SpCas9 Knockout Screens. G3 (Bethesda), 7, 2719–2727.

15. Gonçalves, E., Behan, F.M., Louzada, S., Arnol, D., Stronach, E.A., Yang, F., Yusa, K., Stegle, O., Iorio, F. and Garnett, M.J. (2019) Structural rearrangements generate cell-specific, gene-independent CRISPR-Cas9 loss of fitness effects. Genome Biology, 20, 27.

16. Iorio, F., Behan, F.M., Gonçalves, E., Bhosle, S.G., Chen, E., Shepherd, R., Beaver, C., Ansari, R., Pooley, R., Wilkinson, P., et al. (2018) Unsupervised correction of gene-independent cell responses to CRISPR-Cas9 targeting. BMC Genomics, 19, 604.

17. Langmead, B., Trapnell, C., Pop, M. and Salzberg, S.L. (2009) Ultrafast and memory-efficient alignment of short DNA sequences to the human genome. Genome Biol., 10, R25.

18. Li, W., Xu, H., Xiao, T., Cong, L., Love, M.I., Zhang, F., Irizarry, R.A., Liu, J.S., Brown, M. and Liu, X.S. (2014) MAGeCK enables robust identification of essential genes from genome-scale CRISPR/Cas9 knockout screens. Genome Biol., 15, 554.

19. Ohta, T., Iijima, K., Miyamoto, M., Nakahara, I., Tanaka, H., Ohtsuji, M., Suzuki, T., Kobayashi, A., Yokota, J., Sakiyama, T., et al. (2008) Loss of Keap1 function activates Nrf2 and provides advantages for lung cancer cell growth. Cancer Res., 68, 1303–1309.

20. Taguchi, K. and Yamamoto, M. (2017) The KEAP1-NRF2 System in Cancer. Front Oncol, 7, 85.

21. Couzens, A.L., Knight, J.D.R., Kean, M.J., Teo, G., Weiss, A., Dunham, W.H., Lin, Z.-Y., Bagshaw, R.D., Sicheri, F., Pawson, T., et al. (2013) Protein interaction network of the mammalian Hippo pathway reveals mechanisms of kinase-phosphatase interactions. Sci Signal, 6, rs15.

22. Munoz, D.M., Cassiani, P.J., Li, L., Billy, E., Korn, J.M., Jones, M.D., Golji, J., Ruddy, D.A., Yu, K., McAllister, G., et al. (2016) CRISPR Screens Provide a Comprehensive Assessment of Cancer Vulnerabilities but Generate False-Positive Hits for Highly Amplified Genomic Regions. Cancer Discov, 6, 900–913.

23. Fortin, J.-P., Gascoigne, K.E., Haverty, P.M., Forrest, W.F., Costa, M.R. and Martin, S.E. (2018) Multiple-gene targeting and mismatch tolerance can confound analysis of genome-wide pooled CRISPR screens. bioRxiv, 10.1101/387258.

24. Wienert, B., Wyman, S.K., Richardson, C.D., Yeh, C.D., Akcakaya, P., Porritt, M.J., Morlock, M., Vu, J.T., Kazane, K.R., Watry, H.L., et al. (2019) Unbiased detection of CRISPR off-targets in vivo using DISCOVER-Seq. Science, 364, 286–289.

25. Allen, F., Behan, F., Khodak, A., Iorio, F., Yusa, K., Garnett, M. and Parts, L. (2019) JACKS: joint analysis of CRISPR/Cas9 knockout screens. Genome Res., 29, 464–471.

26. Lahens, N.F., Kavakli, I.H., Zhang, R., Hayer, K., Black, M.B., Dueck, H., Pizarro, A., Kim, J., Irizarry, R., Thomas, R.S., et al. (2014) IVT-seq reveals extreme bias in RNA sequencing. Genome Biol., 15, R86.

27. Parekh, S., Ziegenhain, C., Vieth, B., Enard, W. and Hellmann, I. (2016) The impact of amplification on differential expression analyses by RNA-seq. Sci Rep, 6, 25533.

28. Dempster, J.M., Pacini, C., Pantel, S., Behan, F.M., Green, T., Krill-Burger, J., Beaver, C.M., Younger, S.T., Zhivich, V., Najgebauer, H., et al. (2019) Agreement between two large pan-cancer CRISPR-Cas9 gene dependency data sets. Nat Commun, 10, 5817.

29. Sanson, K.R., Hanna, R.E., Hegde, M., Donovan, K.F., Strand, C., Sullender, M.E., Vaimberg, E.W., Goodale, A., Root, D.E., Piccioni, F., et al. (2018) Optimized libraries for CRISPR-Cas9 genetic screens with multiple modalities. Nat Commun, 9, 1–15.

30. Ghandi, M., Huang, F.W., Jané-Valbuena, J., Kryukov, G.V., Lo, C.C., McDonald, E.R., Barretina, J., Gelfand, E.T., Bielski, C.M., Li, H., et al. (2019) Next-generation characterization of the Cancer Cell Line Encyclopedia. Nature, 569, 503–508.

